# Neonatal diabetes mutations disrupt a chromatin pioneering function that activates the human insulin gene

**DOI:** 10.1101/2020.05.31.125377

**Authors:** Ildem Akerman, Miguel Angel Maestro, Vane Grau, Javier García-Hurtado, Gerhard Mittler, Philippe Ravassard, Lorenzo Piemonti, Jorge Ferrer

## Abstract

Despite the central role of chromosomal context in gene transcription, human noncoding DNA variants are generally studied outside of their endogenous genomic location. This poses major limitations to understand the true consequences of causal regulatory variants. We focused on a cis-regulatory mutation (c.-331C>G) in the *INS* gene promoter that is recurrently mutated in unrelated patients with recessive neonatal diabetes. We created mice in which a ~3.1 kb human *INS* upstream region carrying −331C or −331G alleles replaced the orthologous mouse *Ins2* region. This human sequence drove cell-specific transcription in mice. It also recapitulated poised chromatin during pancreas development and active chromatin in differentiated β-cells. The c.-331C>G mutation, however, blocked active chromatin formation in differentiated b-cells. We further show that another neonatal diabetes gene product, GLIS3, had a singular pioneer-like ability to activate *INS* chromatin in non-pancreatic cells, which was hampered by the c.-331C>G mutation. This *in vivo* analysis of human regulatory defects, therefore, uncovered *cis* and *trans* components of a mechanism that is essential to activate the endogenous *INS* gene.

## INTRODUCTION

Most sequence variants underlying Mendelian diseases affect coding sequences, although a subset of patients harbor cis-regulatory mutations^1–5^. This number is expected to rise as millions of human genomes are sequenced and the field learns to discriminate pathogenic noncoding mutations from a vast number of inconsequential variants^6–9^. In polygenic diseases, common and rare cis-regulatory variants are known to play a central role in the genetic susceptibility^10–12^.

Despite their relevance, cis-regulatory variants have largely been studied outside their true chromosomal context. Current experimental models usually test noncoding variants with episomal DNA constructs, ectopically located transgenes, or *in vitro* protein-DNA interaction assays. The extent to which these models reflect the true impact of cis-regulatory variants is unknown. Quantitative trait loci can provide insights into which genes are affected by regulatory variants in their native genome context, although they do not necessarily model their cellular impact, and generally fail to distinguish causal from linked variants. It is now also possible to directly edit mutations in stem cells and differentiate them *in vitro*, but this does not necessarily allow modeling the mutational impact in relevant developmental or physiological *in vivo* contexts. There is a need, therefore, to develop complementary tools that facilitate understanding of variant pathogenicity and their *in vivo* impact.

One approach to address this need is to engineer human genomic sequences in mice. Several examples of human knock-ins in the mouse genome have been created to model human coding mutations^13^. One study successfully edited a 5 bp noncoding sequence in mice to model a common regulatory variant^14^. However, the extent to which mice can be used to study human cis-regulatory mutations in more extended orthologous genomic contexts is poorly explored.

Modeling noncoding mutations in model organisms poses major challenges. For example, the consequence of a mutation that disrupts a transcription factor-DNA interaction can be influenced by essential combinatorial interactions with nearby bound transcription factors, or by the existence of redundant binding sites^15–17^. Unlike coding sequences, which are often highly conserved, noncoding DNA sequences can maintain functions despite substantial evolutionary turnover, while conserved noncoding sequences can acquire divergent functions^18,19^. The regulatory landscape of individual genes in mouse and human often shares binding sites for ortholog transcription factors, but these are often arranged differently within regulatory elements, and across elements of the same gene. It is thus difficult to predict pathological consequences of human cis-regulatory mutations from mouse models unless the broader sequence context has been humanized.

We have examined a cis-regulatory mutation that causes diabetes mellitus. A subset of patients with neonatal diabetes harbor loss-of-function recessive coding mutations or deletions in *INS*, encoding for insulin^4^. The same phenotype can be caused by *INS* promoter mutations^4^. Interestingly, single nucleotide mutations described so far in >20 probands are located in a CC dinucleotide 331 bp upstream from the *INS* start codon, and −93 bp from the transcriptional start site (c.-331C > G, c.-332C > G, c.-331C > A)^4,20–22^ (**Supplementary Figure 1**). Functional studies in tumoral b-cells using episomal luciferase assays showed that c.-331C>G, the most common of these mutations, causes partial disruption of *INS* promoter activity^4^. However, these mutations have not yet been studied in their *in vivo* genomic context. Decades of work have shown that artificial mutations in numerous elements of the *INS* 5’ flanking region disrupt binding by transcription factors such as MAFA, PDX1,or NEUROD1, and lead to reduced transcriptional activity in episomal assays^23–27^. The reasons why the CC element, rather than other better characterized *INS* promoter functional elements, is the most vulnerable site for loss-of-function single nucleotide mutations in patients remains unknown.

In this study, we generated a mouse model in which the human *INS* upstream genomic region was used to replace the orthologous mouse *Ins2* locus region, and an allelic version that carries the *INS* c.-331C>G mutation. We show that this model recapitulates b-cell specific transcription and acquires expected stage- and cell-specific active chromatin states. Analysis of this *in vivo* model demonstrated that the c.-331C>G mutation does not affect priming of *INS* chromatin during early pancreas development, yet abrogates the formation of active chromatin in differentiated b cells. We further examined GLIS3, a zinc finger transcription factor that is known to regulate the insulin gene^28,29^, and is also mutated in patients with neonatal diabetes and harbors common variants associated with type 1 and type 2 diabetes^30–32^. We show that GLIS3 has a unique capacity among other islet transcription factors to create active chromatin in the *INS* gene in non-pancreatic cells, which is at least partly mediated through interactions with the CC element. Our analysis therefore links two human genetic defects in a mechanism that creates an active chromatin state in the human *INS* gene. These insights are relevant to the genetic mechanisms of diabetes and for regenerative strategies that aim to activate the *INS* gene in non-pancreatic cells.

## RESULTS

### Generation of a human cis-regulatory mutation mouse model

To model the c.-331C>G *INS* mutation in a humanized sequence context, we generated C57BL/6 mice in which homologous recombination was used to replace the endogenous 3.17 Kb mouse genomic region containing the *Ins2* gene and its upstream regions with a human *INS* upstream DNA fragment. This knock-in contained (a) a 3.10 Kb upstream region of human *INS* that included *INS* transcribed 5’ untranslated sequences, (b) mouse *Ins2* exon1 coding sequence, intron, and exon 2, (c) an internal ribosome entry site followed by green fluorescent protein (GFP) sequence (**Figure 1**). The resulting single transcript was thus translated into two proteins: the mouse preproinsulin-2 and GFP. In parallel, we generated a mouse model harboring the same humanized sequence except for the *INS* c.-331C>G single point mutation (**Figure 1**). We named mice carrying the two human *INS* upstream knock-in alleles HIP^KI^ and HIP^KI-*C331G*^.

**Figure 1.**
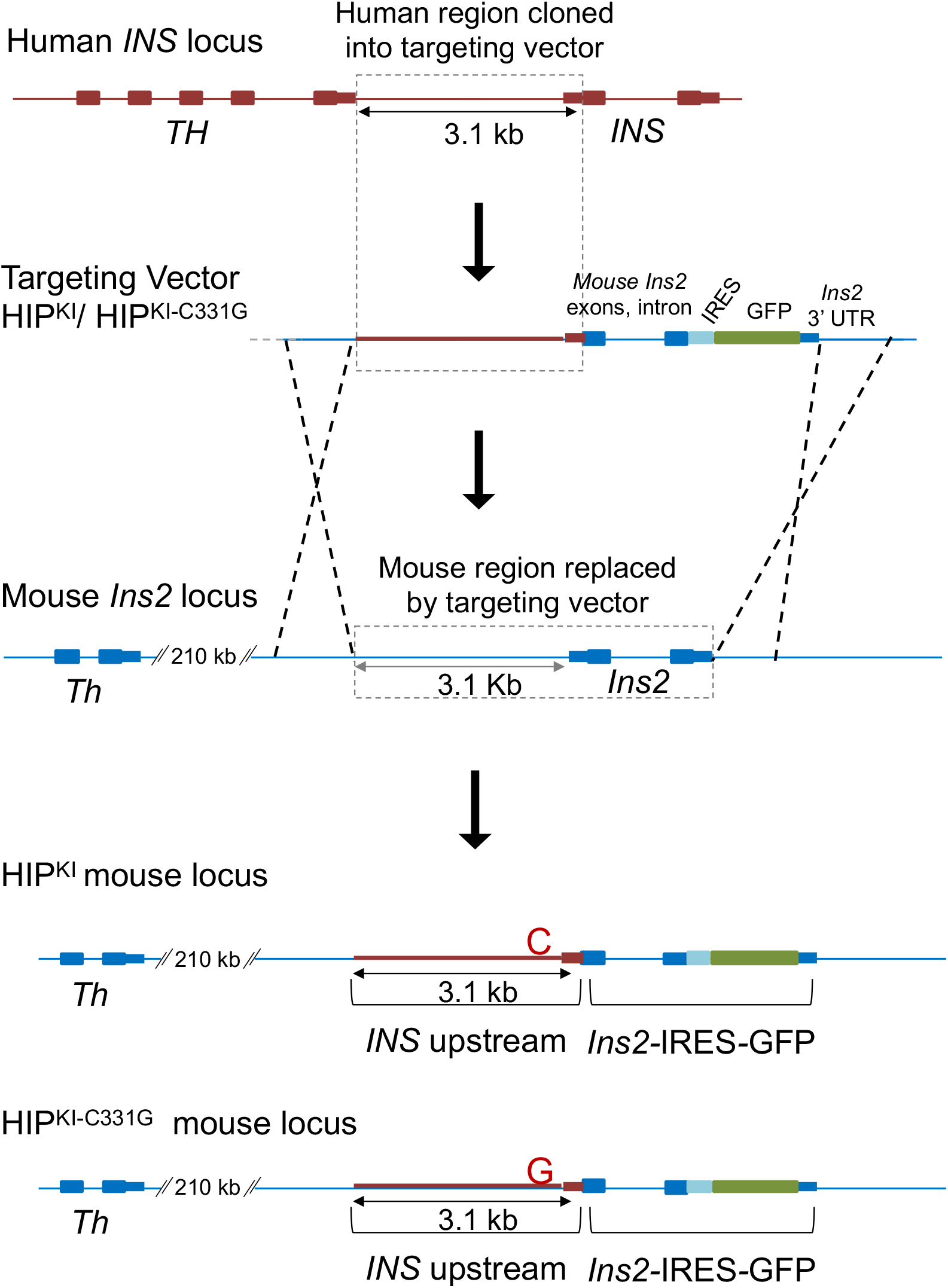
Generation of HIPKI and HIPKI-C331G mouse alleles. A rectangle with dotted lines in the top two panels depicts the 3.1 kb human sequence located between the human *TH* and *INS* genes (including *INS* 5’ untranslated transcribed sequences) which was cloned into a targeting vector. This targeting vector contained the 3.1 kb human INS upstream region followed by Ins2-IRES-GFP, which includes mouse *Ins2* exons and intron, an IRES, GFP, and *Ins2* 3’UTR, and was flanked by mouse *Ins2* homology arms. Targeted replacement of the indicated mouse *Ins2* sequence with this human upstream *INS* sequence followed by Ins2-IRES-GFP was carried out by homologous recombination. The same process was used to create two allelic versions carrying the normal human *INS* sequence or the c.-331C>G mutation. A neomycin cassette flanked by LoxP sites was excised *in vivo*, and is omitted for simplicity. The sequence of the Neomycin-excised targeted allele is provided in Supplementary File 1.

Homozygous HIP^KI^ and HIP^KI-*C331G*^ mice were born at expected Mendelian ratios and appeared healthy without overt hyperglycemia (**Supplementary Figure 2A**), consistent with the fact that HIP knock-in mice retained an intact copy of *Ins1*, a retroposed mouse gene that also encodes insulin^33,34^.

### HIP^KI^ recapitulates, while HIP^KI-*C331G*^ abrogates, cell-specific *INS* transcription

To determine if the human *INS* 5’ flanking region inserted into its orthologous mouse chromosomal context was able to drive b cell-specific expression in HIP^KI^ mice, we used dual immunofluorescence analysis of GFP and islet hormones in tissues from newborn mice (P1-3). This showed that GFP expression was restricted to pancreatic islet core b cells of HIP^KI^ mice, whereas it was not detected in mantle glucagon- or somatostatin-positive islet cells, or surrounding exocrine cells (**Figure 2A**). GFP fluorescence was readily detected in live islets isolated from 3-5 month old HIP^KI^ mice (**Figure 2B**).

**Figure 2.**
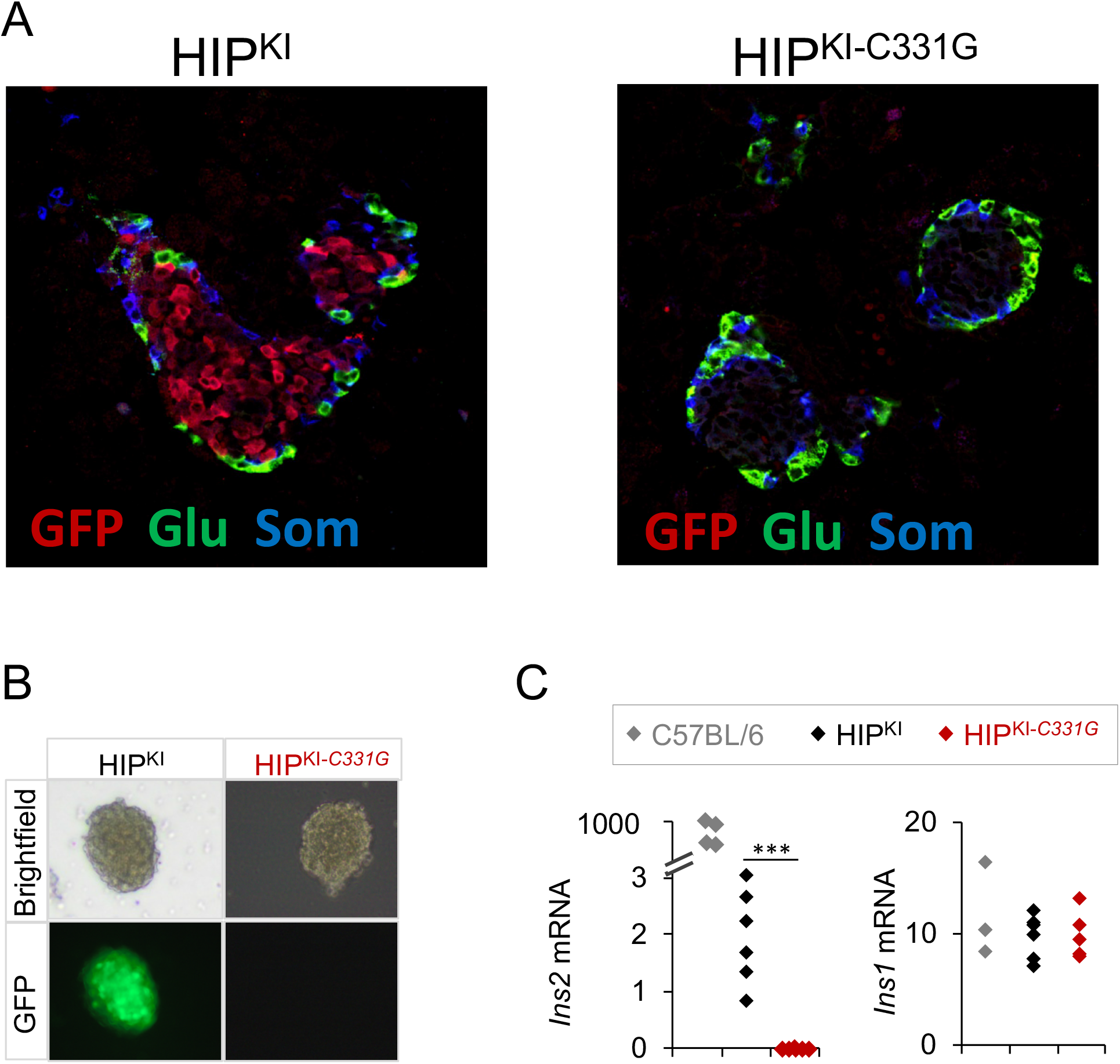
HIP^KI^ recapitulates, and HIP^KI-C331G^ abrogates b cell-specific *insulin* expression. (A) Immunofluorescence imaging of glucagon, somatostatin, and GFP, showing selective GFP expression in HIP^KI^ insulin-positive core islet cells, but not in the surrounding somatostatin or glucagon islet mantle, or in extra-islet exocrine cells, while no GFP expression is detected in HIP^KI-C331G^ islet cells. (B) Brightfield and fluorescence imaging of islets isolated from HIPKI and HIPKI-C331G mice. (C) Quantitative PCR analysis of Ins2 and Ins1 mRNAs in pancreatic islets from 3-5 month old C57BL/6 (n=4), HIP^KI^ (n=6) and HIP^KI-C331G^ (n=6) mice. Values were normalized to Actb mRNA. *** p<0.0001 (Student’s t-test).

In sharp contrast to HIP^KI^ islets, HIP^KI-*C331G*^ islets showed no detectable GFP fluorescence or immunoreactivity (**Figure 2A, B**). Thus, Ins2-IRES-GFP expression in HIP^KI^ mice recapitulates expected β cell-specific patterns of insulin expression, whereas the HIP^KI-*C331G*^ mutation disrupts this expression pattern.

To further assess the function of human *INS* flanking regions, we measured *Ins2* mRNA in islets isolated from HIP^KI^ and HIP^KI-*C331G*^ mice. Quantitative RT-PCR analysis revealed *Ins2* mRNA in islets isolated from both control C57BL/6 and HIP^KI^ mice, but not in HIP^KI-*C331G*^ mouse (**Figure 2C**), thus confirming that the c.-331C>G single point mutation abrogates transcriptional activity of the humanized *INS*/*Ins2* locus in mice. We note, however, that *Ins2* transcripts in HIP^KI^ islets form part of the larger Ins2-IRES-GFP transcript, and were reduced in comparison to control C57BL/6 islets, thus pointing to reduced transcription or stability of *Ins2*-IRES-GFP mRNA (**Figure 2C**)

Further analysis revealed normal levels of *Ins1*, *Pdx1, Glis3, NeuroD1* and *MafA* mRNAs in HIP^KI^ and HIP^KI-*C331G*^ islets, suggesting that humanization of regulatory sequences in the mouse *Ins2* locus did not impact these key pancreatic β-cell identity markers (**Figure 2C, Supplementary Figure 2B**).

Mice with homozygous null *Ins2* mutations display mild transient hyperglycemia^33,34^. Consistently, HIP^KI-*C331G*^ mice, which were *Ins2*-deficient, showed mildly increased glycemia; 119(32) vs. 145(24) mg/dL, mean (IQR), Students t-test P = 0.0031 (**Supplementary Figure 2A**).

These findings indicate that a ~3 kb human *INS* upstream region can replace its mouse orthologous region and still direct cell-specific transcription in mouse b-cells. Furthermore, they show that the human *INS* c.-331C>G point mutation abrogates the function of this humanized region, consistent with the severe phenotype observed in humans with neonatal diabetes.

### HIP^KI^ but not HIP^KI-*C331G*^ islets mirror human islet *INS* chromatin

Nucleosomes that flank promoters of transcriptionally active genes are typically enriched in tri-methylated histone H3 lysine 4 (H3K4me3) and acetylated histone H3 lysine 27 (H3K27ac), and this was expectedly also observed in the human *INS* promoter in human islets^12,35^(**Supplementary Figure 3**). We thus examined these histone modifications at the *INS* promoter in HIP^KI^ and HIP^KI-*C331G*^ adult mouse islets by chromatin immunoprecipitation (ChIP) assays. The *INS* proximal promoter showed enriched H3K4me3 and H3K27ac in adult HIP^KI^ mouse islets (**Figure 3A-C**). By contrast, HIP^KI-*C331G*^ islets showed 5-fold and 7-fold lower signals (Student’s t test, p < 0.001, **Figure 3B,C**). HIP^KI-*C331G*^ islets also showed markedly reduced *INS* promoter chromatin accessibility as measured by FAIRE (**Figure 3D)**, and reduced binding of PDX1, an islet transcription factor that binds to multiple sites in the *INS* promoter (**Figure 3E**). These findings, therefore, demonstrate that the c.-331C>G point mutation does not only prevent transcriptional activity conferred by the human *INS* 5’flanking region but also the formation of accessible active chromatin, and thereby occupancy by a key *INS* gene transcription factor.

**Figure 3.**
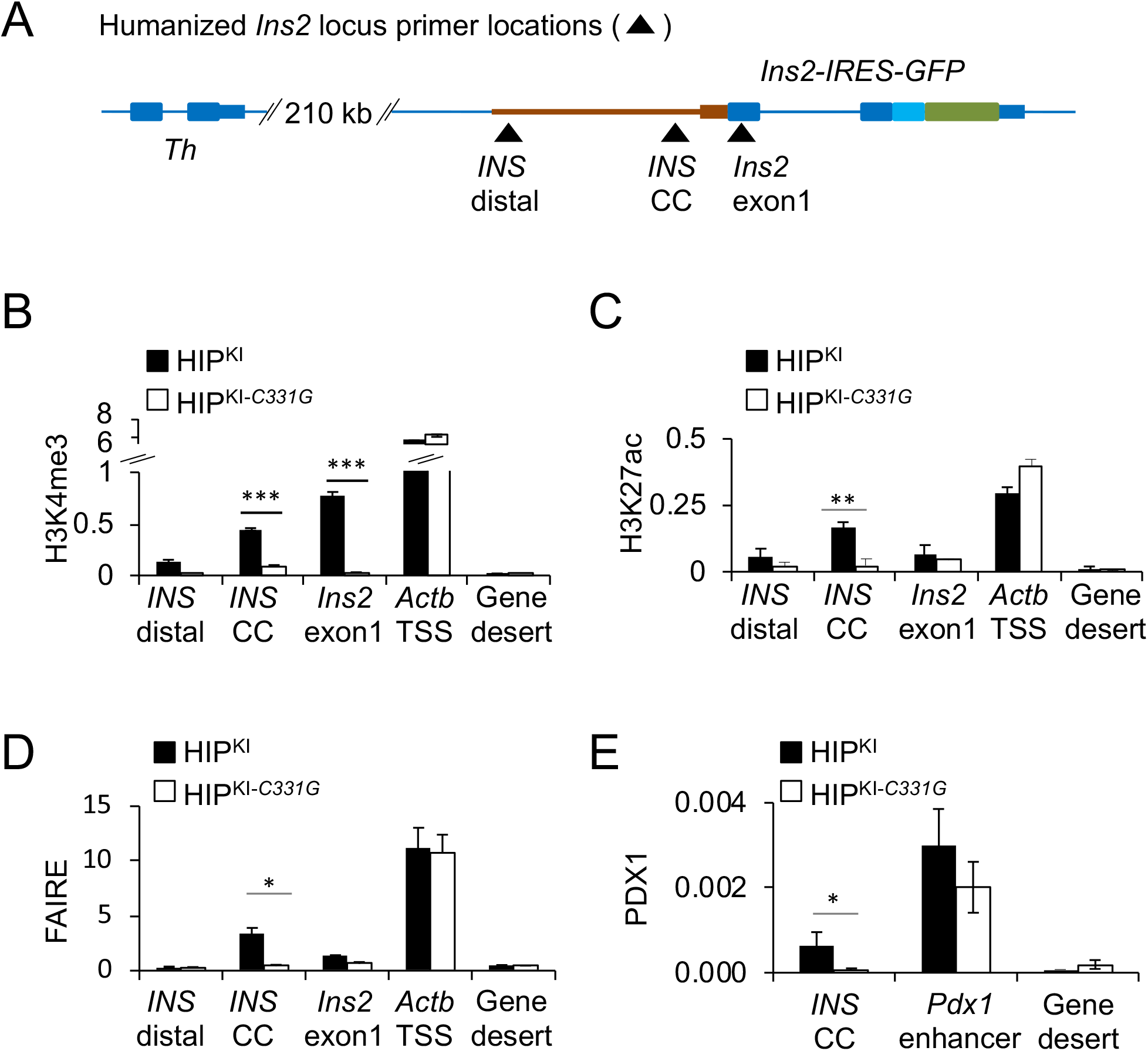
HIP^KI^ recapitulates, and HIP^KI-*C331G*^ abrogates formation of active chromatin at the *INS* gene. (A) Schematic showing approximate primer locations for ChIP assays. *INS* cc encompass the CC element. (B-D) ChIP quantification of H3K4me3, H3K27ac, FAIRE accessibility at the humanized *INS* locus and control regions in HIP^KI^ or HIP^KI-*C331G*^ adult mouse islets. The *Actb* promoter was used as a positive control, and a gene desert lacking active histone modifications in islets was used as a negative control. ChIP and FAIRE DNA were expressed as a percentage of the total input DNA. n= 3 independent experiments. (E) ChIP for the transcription factor PDX1 in HIP^KI^ or HIP^KI-*C331G*^ adult mouse islets at a known PDX1-bound enhancer near *Pdx1* as a positive control. Data is expressed as a percentage of input DNA. n= 3 independent experiments. Error bars indicate SEM, asterisks indicate Student’s t-test P values (* p<0.05, ** p<0.001, *** p<0.0001).

### Activation of humanized *INS* locus during development

We next used HIP^KI^ and HIP^KI-*C331G*^ models to assess how the c.-331C>G mutation influences chromatin activation at the *INS* gene during pancreas development. As a reference, we analyzed ChIP-seq maps of H3K4me1 and H3K4me3 histone modifications in human fetal pancreas at Carnegie stage 23^36^, which precedes *INS* gene activation, and compared them with analogous maps from adult human islets^12,35^. In the human fetal pancreas, we observed that the histone modification H3K4me1 demarcated a broad region that encompasses the *TH* and *INS* genes as well as downstream regions, without detectable H3K4me3, a histone modification associated with active promoters (**Figure 4A)**. This combination, H3K4me1 enrichment and absence of H3K4me3, has been described in genomic regions that are poised for gene activation^37^, and is consistent with the virtual absence of *INS* transcription in the early fetal pancreas. In adult human islets, by contrast, H3K4me1 in the *INS* locus was largely replaced with H3K4me3, in keeping with active *INS* transcription (**Figure 4A**).

**Figure 4.**
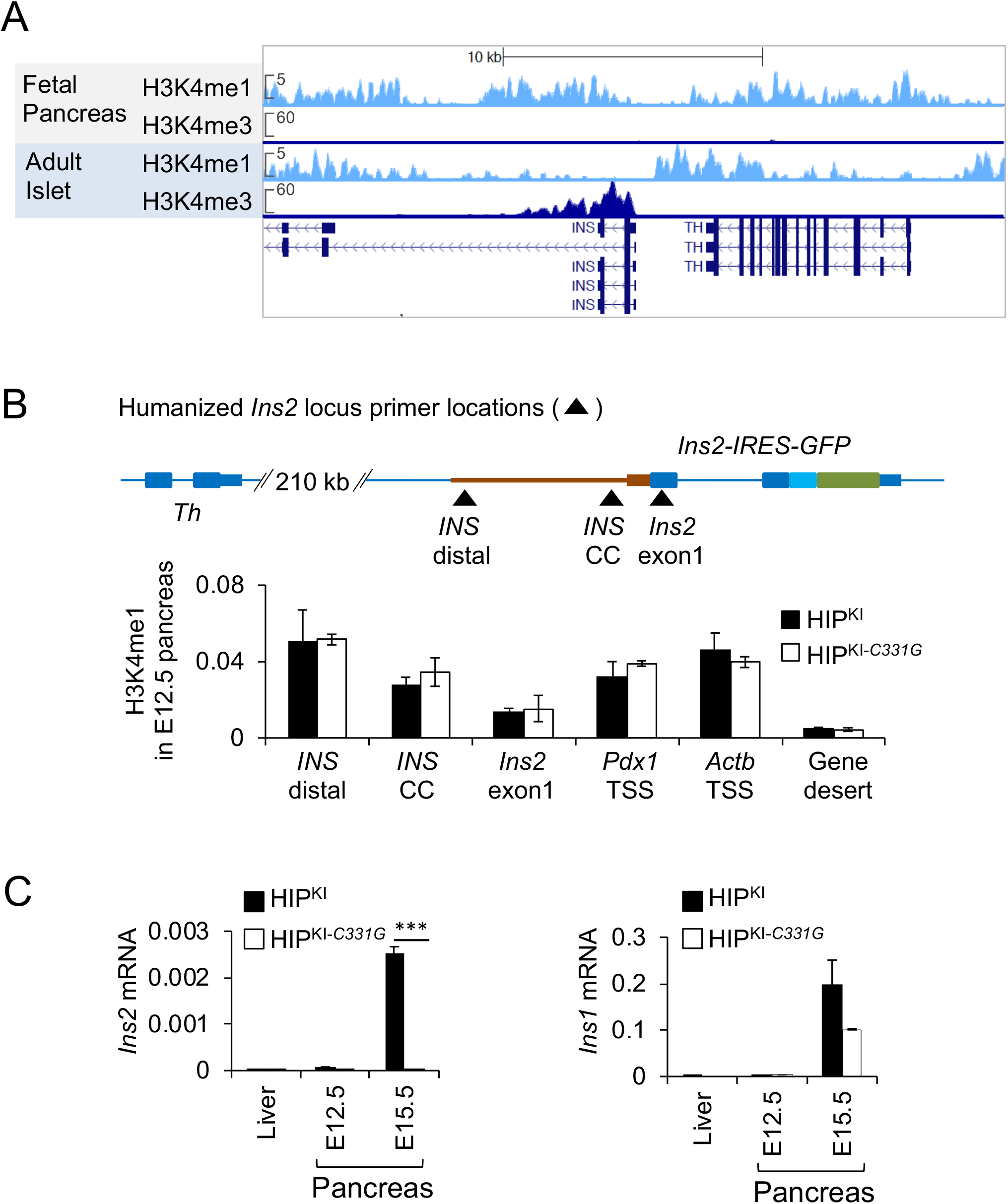
Developmental activation of *INS* promoter chromatin. (*A*) H3K4me1 and H3K4me3 profiles at the *INS* locus showing H3K4me1, but not H3K4me3, enrichment in human fetal pancreas (Carnegie stage 23)^36^ across the *INS* region, and H3K4me3, but not H3K4me1, enrichment in adult human islets^12,35^. (B) Top panel: approximate location of oligonucleotides for ChIP assays. Bottom panel: ChIP H3K4me1 enrichments in pancreas from HIP^KI^ or HIP^KI-*C331G*^ E12.5 embryos. N = 2 independent experiments from pools of 8-12 embryos. Data is expressed as a percentage of input DNA. **(C)** Quantitative PCR analysis for *Ins2* and *Ins1* mRNA from HIP^KI^ or HIP^KI-*C331G*^ E12.5 and E15.5 embryonic pancreas or E15.5 liver (n=3). Error bars indicate SEM, asterisks indicate Student’s t-test P values *** p<0.0001.

We next examined if the chromatin environment that precedes transcriptional activation of the *INS* locus is recapitulated in HIP^KI^ embryos. As in human fetal pancreas, we observed deposition of H3K4me1 in the humanized *INS* flanking regions in HIP^KI^ E12.5 mouse fetal pancreas (**Figure 4B**). *Ins2* mRNA was expectedly not detected above background levels at this stage (**Figure 4C**). Interestingly, both HIP^KI^ and HIP^KI-*C331G*^ displayed similar H3K4me1 profiles at E12.5 (**Figure 4B**), indicating that the humanized *INS* sequences recapitulate poised chromatin preceding gene activity, and this is not impeded by the c.-331C>G mutation.

By contrast, the c.-331C>G mutation prevented the transcriptional activation of the humanized locus at early stages of b cell differentiation. Thus, *Ins2* mRNA was detectable in *HIP*^KI^ E15.5 fetal pancreas, but it was undetectable in HIP^KI-*C331G*^ embryos (**Figure 4C**), whereas *Ins1* mRNA was readily detected in both HIP^KI^ and HIP^KI-*C331G*^ E15.5 pancreas (**Figure 4C**). Thus, mice carrying humanized regulatory regions in the *Ins2* locus can recapitulate salient developmental features of the chromatin landscape of the human *INS* locus, and show that although the c.-331C>G mutation does not affect *INS* chromatin poising, it disrupts activation of promoter chromatin in differentiated cells.

### GLIS3-dependent activation of the *INS* gene is prevented by c.-331C>G

We next sought to identify factors whose DNA binding activity is influenced by the c.-331C>G mutation. We performed SILAC experiments, and identified four zinc finger transcription factors (MAZ, ZFP37, KLF13, KLF16) that showed decreased *in vitro* binding to the c.-331C>G mutation, and additionally selected 5 zinc finger transcription factors expressed in human islets that were predicted to show differential binding based on *in silico* and/or literature analysis (**Supplementary Figure 4A**). Of these 9 candidate transcription factors, GLIS3, which was previously shown to bind to this element *in vitro*^28^, induced significant luciferase activity in a human insulin promoter episomal construct, while the c.-331C>G mutation suppressed this effect (T-test p = 0.02, **Figure 5A**). Systematic DNA binding site selection studies predict that the C>G mutation impairs GLIS3 binding^38^, and this was confirmed with electromobility shift assays (**Supplementary Figure 4B**). These findings, therefore, indicated that the c.-331C>G mutation disrupts GLIS3 *in vitro* binding to the CC element as well as activation of an episomal *INS* promoter.

**Figure 5.**
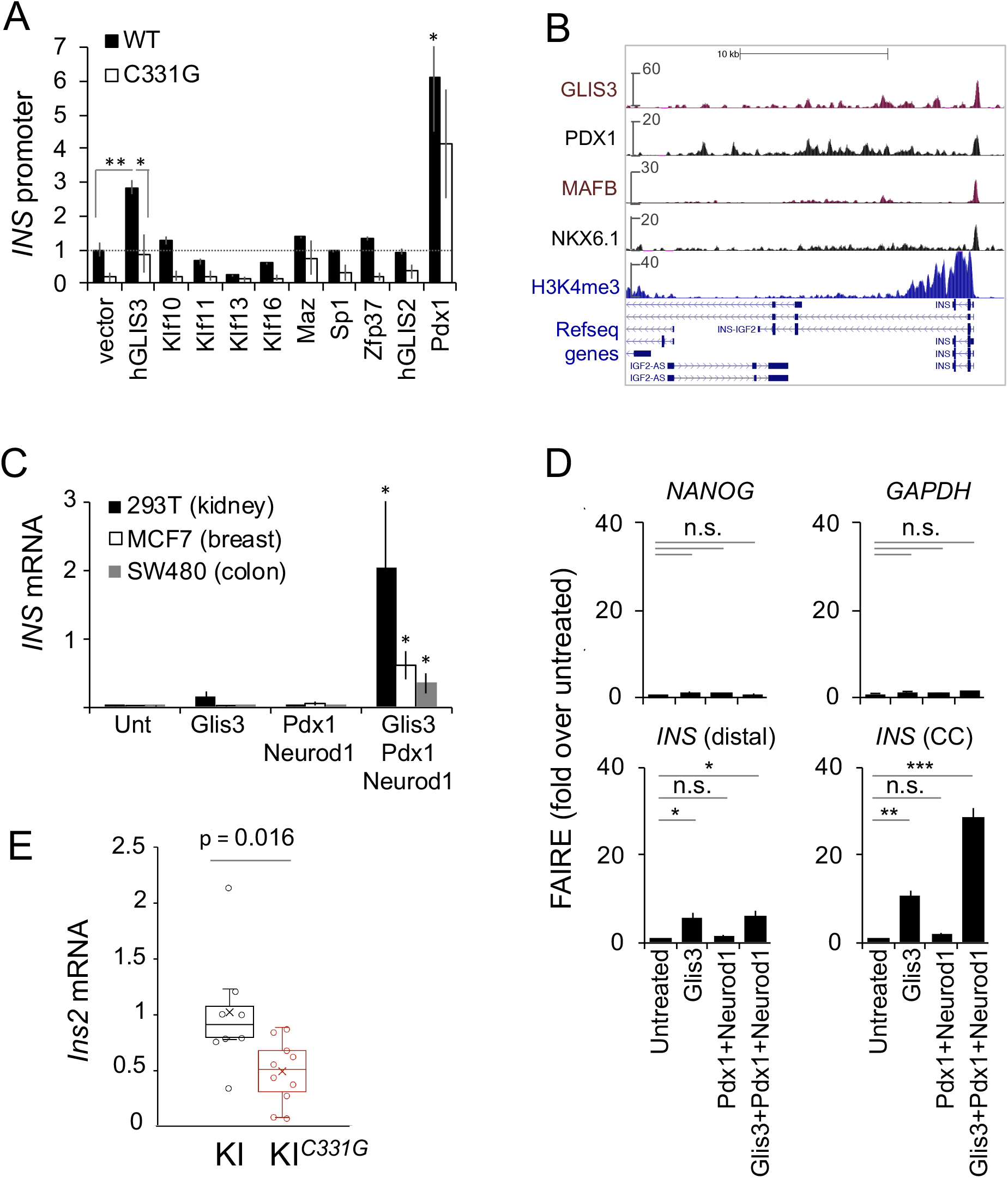
GLIS3 regulates activates the endogenous *INS* gene and requires an intact CC element. (A) Transfection of candidate transcription factors or PDX1 as a positive control, along with INS promoter (wild type or c.-331C>G) luciferase and control renilla plasmids in ENDOCβ-H1 cells. Statistical comparisons are with an empty expression vector. For hGLIS3 a comparison of WT vs. c.-331C>G is also shown. (B) ChiP-seq shows GLIS3 binding to the *INS* promoter in human islets. (C) GLIS3 activates *INS* in heterologous cell types. HEK 293T, MCF7 or SW480 cells were transfected with plasmids encoding indicated transcription factors or left untreated (unt). *INS* mRNA was calculated as *INS* to *GAPDH* mRNA ratios X 1000. Significance was calculated relative to untreated samples in 3 independent experiments. (D) FAIRE assessment of accessible chromatin in transfected HEK 293T cells. FAIRE DNA was quantified by PCR and expressed as percentage of input DNA and fold enrichment over untreated cells. Statistical significance was calculated relative to untreated samples (n=3 independent experiments). Expectedly, *NANOG* and *GAPDH* did not change. (E) GLIS3-dependent activation of *INS* in fibroblasts obtained from HIP^KI^ (n=7) and HIP^KI-C331G^ (n=10) embryos. Fibroblasts were transduced with mouse GLIS3, MAFA, PDX1 and NEUROD1 lentivirus, and RNA was analyzed after 2 days. Two independent experiments were performed with cells from 7 HIP^KI^ and 10 HIP^KI-C331G^ embryos from two litters. Error bars are SEM, asterisks are Student’s t-test p<0.05 (*), p<0.001 (**), p<0.0001(***).

Recently, GLIS3 was shown to bind *in vivo* to the *Ins2* promoter, while pancreatic inactivation of *Glis3* in mice specifically depletes *Ins2* mRNA in islet cells^39,40^. To test whether GLIS3 is a direct *in vivo* regulator of the human *INS* gene, we performed chromatin immunoprecipitation coupled to sequencing (ChIP-seq) of GLIS3 in human islets. This showed 746 high-confidence binding sites, which displayed marked enrichment of canonical GLIS3 binding sequences, including a *de novo* motif matching the *INS* promoter CC element, and a specific binding site in the *INS* promoter (**Figure 5B, Supplementary Figure 5A, Supplementary Table 1**). Additional GLIS3-bound regions were observed in genes known to be important for islet cells including *PDX1*, *MAFA*, *CREB1*, *DLL1* (**Supplementary Figure 5B**). Furthermore, transduction of two independent shRNAs that reduced *GLIS3* mRNA in human EndoCb-H1 b-cells led to ~2-fold lower *INS* mRNA levels (t-test p < 0.05, **Supplementary Figure 5C**). This provided *in vivo* evidence that GLIS3 is a direct regulator of the human *INS* gene.

To understand GLIS3-mediated regulation of *INS* locus chromatin, we assessed the ability of GLIS3 to activate *INS* in three immortalized non-pancreatic human cell lines in which the *INS* gene is repressed. Combinations of islet cell transcription factors, such as PDX1, MAFA, and NEUROG3 or NEUROD1, have been used to activate beta cell programs in pancreatic acinar or liver cells^41,42^, in line with the knowledge that PDX1, MAFA, and NEUROD1 directly bind and regulate the insulin gene promoter in various species^27,43^. We found that expression of various combinations of such transcription factors failed to elicit major changes in *INS* mRNA in cell lines from distant lineages, namely HEK 293T, MCF7 or SW480 cells (**Figure 5C, Supplementary Figure 6A**). By contrast, co-transfection of these factors with GLIS3 led to ectopic activation of the endogenous *INS* gene (**Figure 5C, Supplementary Figure 6A**). Other candidate transcription factors that were predicted to bind to the CC element failed to activate *INS* in the presence of PDX1, MAFA, and NEUROD1 or NEUROG3 (**Supplementary Figures 4A and 6A**). Microarray analysis of HEK293T cells transfected with GLIS3, PDX1, NEUROD1 and MAFA showed that *INS* was the most highly induced gene relative to cells that were only transfected with PDX1, MAFA and NEUROD1. (**Supplementary Figure 6B**). These results indicate that GLIS3 has a specific ability to activate the *INS* gene in cell lines from distant lineages, which requires other *INS* promoter-binding transcription factors that do not elicit this effect on their own.

To further understand this singular effect of GLIS3 on the *INS* gene, we examined chromatin accessibility using FAIRE. Misexpression of GLIS3, PDX1 and NEUROD1 created accessible chromatin at the endogenous *INS* gene in HEK293T cells, whereas this was not elicited by PDX1 and NEUROD1 alone (**Figure 5D).** This effect was direct, as ChIP experiments showed that transfected GLIS3 was bound to the *INS* promoter **(Supplementary Figure 6C**). Thus, GLIS3 has a singular ability to bind and activate the *INS* promoter in repressed chromatin cellular environments.

Because GLIS3 activation of episomal INS promoter constructs were impaired by the c.-331C>G mutation, we next tested whether the GLIS3-dependent chromatin pioneering function was also mediated by the CC element. We thus prepared fibroblasts from HIP^KI^ and HIP^KI-*C331G*^ embryos. Given that other islet transcription factors are required for the GLIS3 effect, we transduced HIP^KI^ and HIP^KI-*C331G*^ mouse embryonic fibroblasts with PDX1, NEUROD1, MAFA and GLIS3. This combination of transcription factors activated transcription from the humanized locus in HIP fibroblasts, whereas the effect was inhibited by the c.-331C>G mutation (**Figure 5E**).

These findings, therefore, indicate that GLIS3 has a unique pioneering function in the activation of human *INS* in an endogenous chromosomal context, and indicate that c.-331C>G mutation impairs this function.

## DISCUSSION

We have used *in vivo* approaches to understand the consequences of two genetic defects that cause human neonatal diabetes. We humanized a large genomic regulatory region in mice, and demonstrated that point mutations that cause neonatal diabetes disrupt an essential pioneering step in the activation of the human *INS* gene. We further examined GLIS3, encoded by another gene that is mutated in neonatal diabetes and harbors type 2 and type 1 diabetes risk variants^30–32^. This showed that GLIS3 also has a unique role in the activation of *INS* gene chromatin, that is at least partly mediated by the same *INS* promoter sequence that is mutated in neonatal diabetes. These results revealed *cis* and *trans* regulators of a previously unrecognized mechanism that activates the endogenous *INS* gene.

Transcriptional regulatory DNA variants play a central role in human disease^44^, yet so far most efforts have investigated their function in experimental systems that do not consider their *in vivo* impact^45–47^. This is a major limitation, because it is currently clear that gene regulation entails a complex interplay between elements that are difficult to reproduce outside of an *in vivo* context, including chromatin structure, epigenetic chemical modifications, or noncoding RNAs. Such factors are highly dynamic throughout development and physiological settings. Importantly, *in vitro* models cannot easily examine transcription factor-DNA interactions that overcome repressed chromatin states. Many transcription factors can only bind recognition sequences in accessible DNA, whereas a subset of transcription factors have the ability to bind to nucleosomal-bound DNA and to reprogram silent chromatin^48,49^. Such pioneer functions play a major role in differentiation and cellular programming, and can thus be studied with *in vivo* models.

Previous work with a trans-species aneuploid model showed that the human chromosome 21 can recapitulate human cell-specific regulatory landscapes in mice^50^. Our study has now shown that a human ~3.1 kb genomic region could be integrated into an orthologous region of another mammal and recapitulate stage and cell-specific functions. To our knowledge, this is the first study that has modeled a human noncoding mutation in an extended human regulatory sequence integrated in an orthologous mouse locus^13^. All point mutations of the *INS* promoter identified so far, including the c.-331C>G mutation, fall in the same CC element. Human genetics thus points to a unique role of this dinucleotide sequence. Our experiments showed that the c.-331C>G mutation did not prevent *INS* chromatin priming in predifferentiated cells, but disrupted H3K4 trimethylation, chromatin accessibility and transcription factor binding to the *INS* promoter in differentiated b-cells. This indicates that the CC element acts as an essential seeding site for chromatin opening and transcriptional activation of the *INS* promoter in b-cell development.

Our studies also show that the Krüppel-like zinc finger protein GLIS3 activates the human *INS* gene *in vivo*. This extends studies showing that GLIS3 binds the mouse *Ins2 in vivo* and the human *INS* CC element *in vitro*^28,51^, while pancreatic *Glis3*-deficient and *Glis3*^+/-^ mice show severe abnormalities in insulin expression^39,40^. More importantly, our studies now show that GLIS3 has a selective ability amongst known *INS* gene regulators to activate *INS* in silent chromatin cellular environments. It is interesting to note that GLIS1, a GLIS3 paralog, has a major impact on reprogramming of pluripotent cells from somatic cells in the presence of other pluripotency transcription factors^52^, while GLIS3 has a similar reprogramming function in some somatic cell lineages^53^. It is therefore likely that GLIS3 can derepress diverse target genes through combinatorial interactions with cell-specific transcription factors. We have further shown that the c.-331C>G mutation not only prevented GLIS3-dependent activation of an episomal *INS* promoter, but also impaired activation of the endogenous genes. These findings, therefore, provide a common mechanism for *GLIS3* and the *INS* CC element genetic defects.

Interestingly, systematic *in vitro* binding studies and episomal reporter assays have disclosed >16 binding activities and functional elements in the human insulin promoter, yet did not highlight the critical role of the CC sequence prior to the human genetic findings^4,22–26^. A mutation of a 34 bp region that contains the CC element in randomly integrated human *INS* promoter transgenics did not significantly alter promoter activity^54^. This indicates that the essential function of the CC element to create active chromatin can only become fully apparent in a natural chromatinized environment, such as that of human patients or mutant HIP mice.

Our studies, therefore, uncover an unanticipated protagonism of GLIS3 and the CC element of the *INS* promoter to initiate an active chromatin state at the human *INS* gene. Our findings are relevant to understanding genetic mechanisms underlying diabetes, as well as for efforts to use *cis*-acting sequences and transcription factors to activate b cell genes replacement therapies^55–57^.

## MATERIALS AND METHODS

### Generation of HIP^KI^ and HIP^KI-*C331G*^ mice

Mouse experiments were conducted following procedures approved by the Ethical Committee of Animal Experimentation of the University of Barcelona. Targeted replacement of *Ins2* with human *INS* 5’ flanking sequences driving *Ins2* and GFP was performed as schematized in **Figure 1** (genOway). The exact sequences of these regions are provided in **Supplementary File 1**. In brief, targeting vectors were generated with a 3.10 kb unmodified or mutated (c.C331G) 5’ flanking *INS* region, *Ins2* exon 1, intron, and exon 2, an IRES-GFP reporter cassette inserted in the 3’ untranslated region of *Ins2* exon 2, followed by a neomycin selection cassette flanked by loxP sites. This construct was flanked by 5.3 kb and 1.8 kb C57BL/6J mouse homology arms. Targeting of this vector was carried out by homologous recombination in C57BL/6J embryonic stem cells, verified by PCR screening and southern blotting. Four suitable clones resulted, which were used for blastocyst injections and generation of the HIP knock-in mouse strains.-This was followed by CRE-mediated excision of the neomycin cassette in vivo, and again verified by PCR screening and southern blotting. The sequence of the replaced region was verified by Sanger sequencing. Oligonucleotides used for genotyping are shown in **Supplementary Table 2**.

### Immunofluorescence

Pancreases were processed for immunofluorescence as previously described^58^. Briefly, tissues were fixed in 4% paraformaldehyde overnight at 4°C, then washed in PBS before paraffin embedding. 4 μm sections were deparaffinized in xylene and rehydrated with ethanol series. Sections were incubated for 30 min at room temperature in antibody diluent (DAKO Corporation) with 3% normal serum from the same species as the secondary antibody and incubated overnight at 4 °C with primary antibodies, then overnight at 4 °C with secondary antibody. Images were acquired using Leica TSE confocal microscope for immunofluorescence. Antibodies used are shown in **Supplementary Table 2**.

### Mouse islets and embryonic pancreas

Mouse islets were isolated using previously described protocols^59^. Unless otherwise indicated, islets were incubated in 4 mM glucose RPMI 1640 medium, supplemented with 10% fetal calf serum, 100 U/ml penicillin, and 100 U/ml streptomycin for 48 hours after isolation. For ChIP experiments, mouse islets were immediately crosslinked after isolation with formaldehyde and frozen until use. Pancreas were dissected from timed mouse embryos and processed as described^60^.

### Human islets

Human pancreatic islets were obtained through the European Consortium on Islet Transplantation (ECIT), Islets for Basic Research Program supported by the Juvenile Diabetes Research Foundation (program 2-RSC-2019-724-I-X). Pancreatic islets were isolated from multiorgan donors without a history of glucose intolerance^61^, shipped in culture medium and re-cultured at 37°C in a humidified chamber with 5% CO_2_ in glucose-free RPMI 1640 supplemented with 10% fetal calf serum, 100 U/ml penicillin, 100 U/ml streptomycin and 11mM glucose for three days before analysis.

### FAIRE, ChIP and ChIP-Seq

ChIP, ChIP-seq and FAIRE (Formaldehyde-Assisted Isolation of Regulatory Elements) were performed using previously published protocols^12,62,63^. H3K4me1 and H3K4me3 ChiPs were performed on ~250 islet equivalents (IEQ) per sample, and PDX1 ChIP experiments were performed on ~400 IEQ per sample. FAIRE experiments were performed on ~250 IEQ, isolated from multiple mice using previously published protocols^62^.

GLIS3 ChiP-seq experiments were performed with ~3000 human IEQ, using a previously described antibody^64^(**Supplementary Table 2**). Sequencing and processing were performed as described^12^. In brief, reads were aligned to the human genome (hg19) using bowtie (default options). Peaks were called using MACS2 (q-value cutoff 0.05, band width 300) against a Input DNA bam file. GLIS3-bound regions are provided in **Supplementary Table 1.**

Antibodies and oligonucleotides are provided in **Supplementary Table 2**. References for new and previously reported human islet and fetal pancreas CHIP-seq datasets used here^12,35,36^ are provided in **Supplementary Table 3**.

### In silico motif analysis

To identify candidate transcription factors that bind the CC element, we performed in silico motif searches using the MEME suite^65^ using the normal and mutated (c.331 C>G) sequences. Analysis of de novo and previously characterized position weight matrixes (PWM) was performed with HOMER v3.12^66^ using peak summit coordinates and flags −size −50,50 −len 6,8,10,12.

### PCR analysis of mRNAs

RNA was isolated using TriPure reagent as described^67^. *Ins2*, *INS* and ActB transcripts were measured using TaqMan (Applied Biosystems) probes, while other regions were quantified using oligonucleotide primers and SYBR green mastermix (Illumina). All mRNA levels were normalized to *ActB* or *Hprt* as indicated in Figure legends. Oligonucleotides can be found in **Supplementary Table 2**.

### GLIS3 knockdown in EndoCb-H1 cells

EndoCβ-H1 cells were cultured as described^68^. Lentiviral-mediated knockdown of GLIS3 in the human pancreatic β-cell line EndoCb-H1 was performed with two independent shRNAs placed in artificial miRNAs as described^67^. Oligonucleotides used for shRNAs are shown in **Supplementary Table 2**.

### Luciferase assays

*INS* promoter activity was measured using pSOUAPRL-251hINS-Luc (Roland Stein, Vanderbilt University), which contains 0-251 bp of human *INS* proximal promoter DNA located in front of the Firefly luciferase cDNA. The c.C331G mutation was introduced using site directed mutagenesis^4^. The plasmids were transfected in in EndoCb-H1 with Lipofectamine 2000 at 4:1 luciferase:over-expression vector ratio as previously described^69^. We co-transfected a Renilla expressing construct (pGL4.75, 0.02ng) as a normalizer to correct for differences in transfection efficiency. Expression vectors used in co-transfections are given in **Supplementary Table 2.**

### Activation of endogenous *INS* in heterologous cell types

Mouse embryonic fibroblasts (MEFs) were obtained from E15.5 HIP^KI^ and HIP^KI-*C331G*^ mouse embryos by clipping the tail tips and culturing in media. HIP^KI^ and HIP^KI-*C331G*^ MEFs, HEK 293T, MCF7 and SW480 cells were maintained in DMEM (Lonza) and supplemented with 10% fetal calf serum, 100 U/ml penicillin, 100 U/ml streptomycin at 37°C in a humidified chamber with 5% CO2. Embryonic fibroblasts cells were transduced with lentiviral vectors, one encoding the transcription factors PDX1, NEUROD1 and MAFA in a polycistronic transcript, and the other GLIS3-ΔN155^38^ (**Supplementary Table 2**). Cells were harvested 48 hours post transduction and subjected to quantitative PCR analysis. HEK 293T, MCF7 and SW480 cells were transfected using Lipofectamine 2000, following manufacturer’s instructions. We note that the omission of MAFA or substitution of NEUROD1 with NEUROG3 in such experiments resulted in similar *INS* induction levels. For microarray hybridization experiments described in **Supplementary Figure 6**, harvested RNA was hybridized to Gene ST 1.0 Affymetrix arrays and the data was analyzed on Affymetrix TAC software as described^12^.

### SILAC

We used Stable Isotope Labeling by Amino acids in Cell culture (SILAC) for mass spectrometry identification of proteins that specifically bound to unmodified or mutated (c.C331G) double stranded oligonucleotides containing a 5’ linker and PstI restriction site (**Supplementary Table 2**), exactly as described^70^, using MIN6 immortalized mouse b cells^71^. This identified 4 proteins whose binding was affected by the mutation: Krueppel-like factor 13 (UniProtKB/Swiss-Prot Q9Y2Y9), Krueppel-like factor 16 (UniProtKB/Swiss-Prot Q9BXK1), MAZ (UniProtKB/TrEMBL Q8IUI2) and ZFP37 (UniProtKB/Swiss-Prot Q9Y6Q3).

## Acknowledgements

This research was supported by the Birmingham Fellowship Programme to I.A. and grants to J.F. from Ministerio de Ciencia e Innovación (BFU2014-54284-R, RTI2018-095666-B-I00), Medical Research Council (MR/L02036X/1), Wellcome Trust (WT101033), European Research Council Advanced Grant (789055), and FP6-LIFESCIHEALTH 518153. Human islets for research were supported by the Juvenile Diabetes Research Foundation (2-RSC-2019-724-I-X). Work in CRG was supported by the CERCA Programme, Generalitat de Catalunya and Centro de Excelencia Severo Ochoa (SEV-2015-0510). We thank the University of Barcelona School of Medicine animal facility, Center of Genomic Regulation and Imperial College London Genomics Units, and also Larry Chan (Baylor College), Roland Stein (Vanderbilt University), Anton Jetten (NIEHS, NIH, North Carolina), Doris Stoffers (University of Pennsylvania), Jochen Seufert (University of Freiburg), Marko Horb (Marine Biological Laboratory), Alpana Ray (University of Missouri), Tatsuya Tsurimi ( Aichi Cancer Center, Ngoya, Japan) for generous gifts of valuable reagents. We thank Diego Balboa and Mirabai Cuenca for critical comments on the manuscript.

## Author Contributions

I.A., J.F. conceived and coordinated the study. I.A. performed cell based and computational studies, supervised mouse analysis. I.A., M.A.M. performed image analysis of mouse mutants and isolated islets. L.P. purified human islets. V.G. maintained mouse colonies, G.M. performed SILAC experiments, P.R. designed and created lentiviral vectors, I.A., J.F. wrote the manuscript with input from the remaining authors.

## Data availability

GEO accession number for GLIS3 ChIP-seq data: GSE151405.

## Figure legends

**Supplementary Figure 1.**
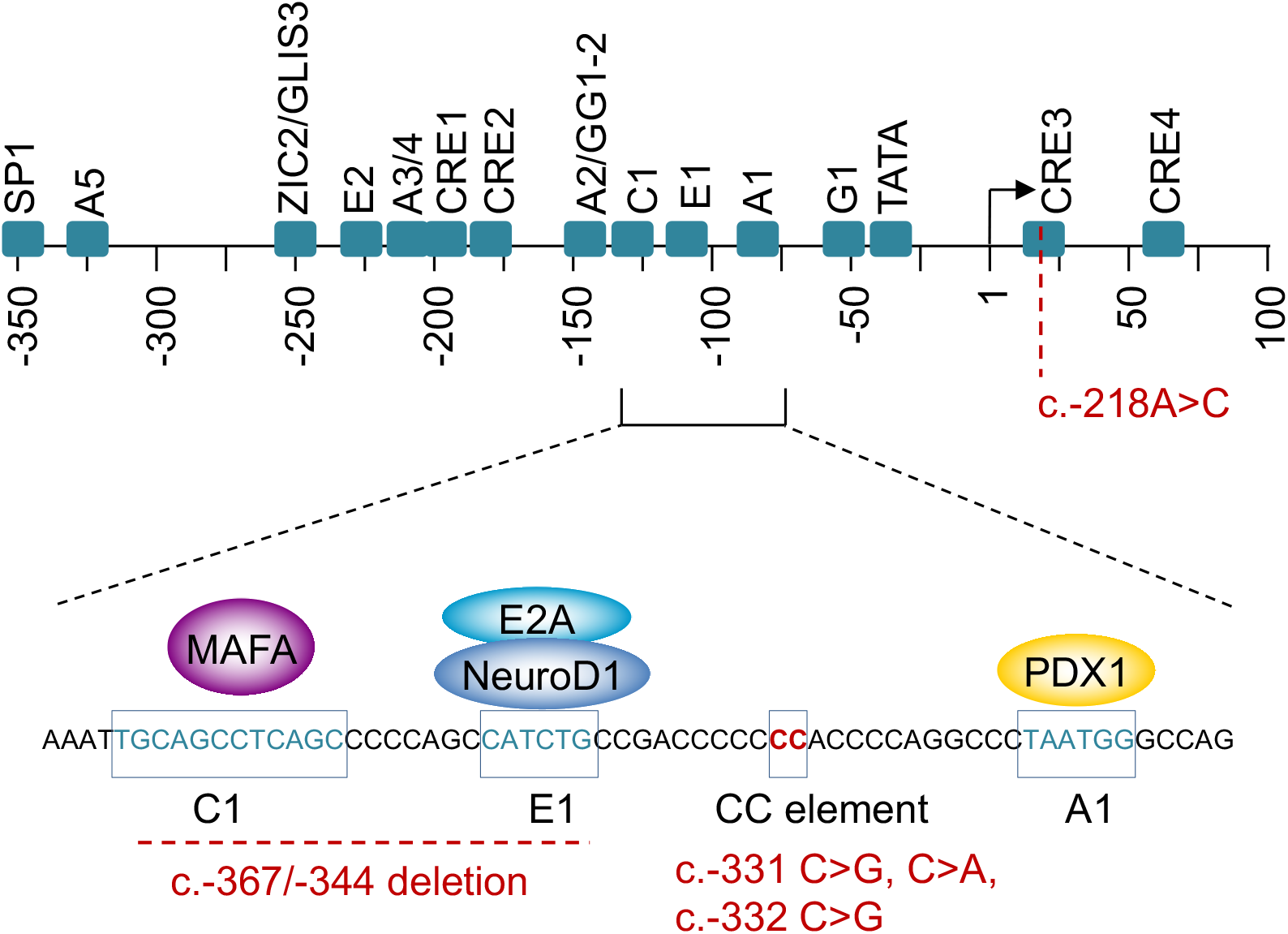
Schematic of the human *INS* 5’ flanking sequence, showing approximate locations of established cis-regulatory sequences based on mutational analysis of episomal sequences. The bottom panel shows a zoomed in sequence that contains MAFA, NEUROD1, and PDX1-bound cis-elements, as well as CC element. Recessive mutations in patients with neonatal diabetes are shown in red. These include 3 point mutations in the CC element, a 17 bp deletion that disrupts MAFA and NEUROD1 binding sites, and a point mutation in the 5’ untranslated INS mRNA sequence^4^.

**Supplementary Figure 2.**
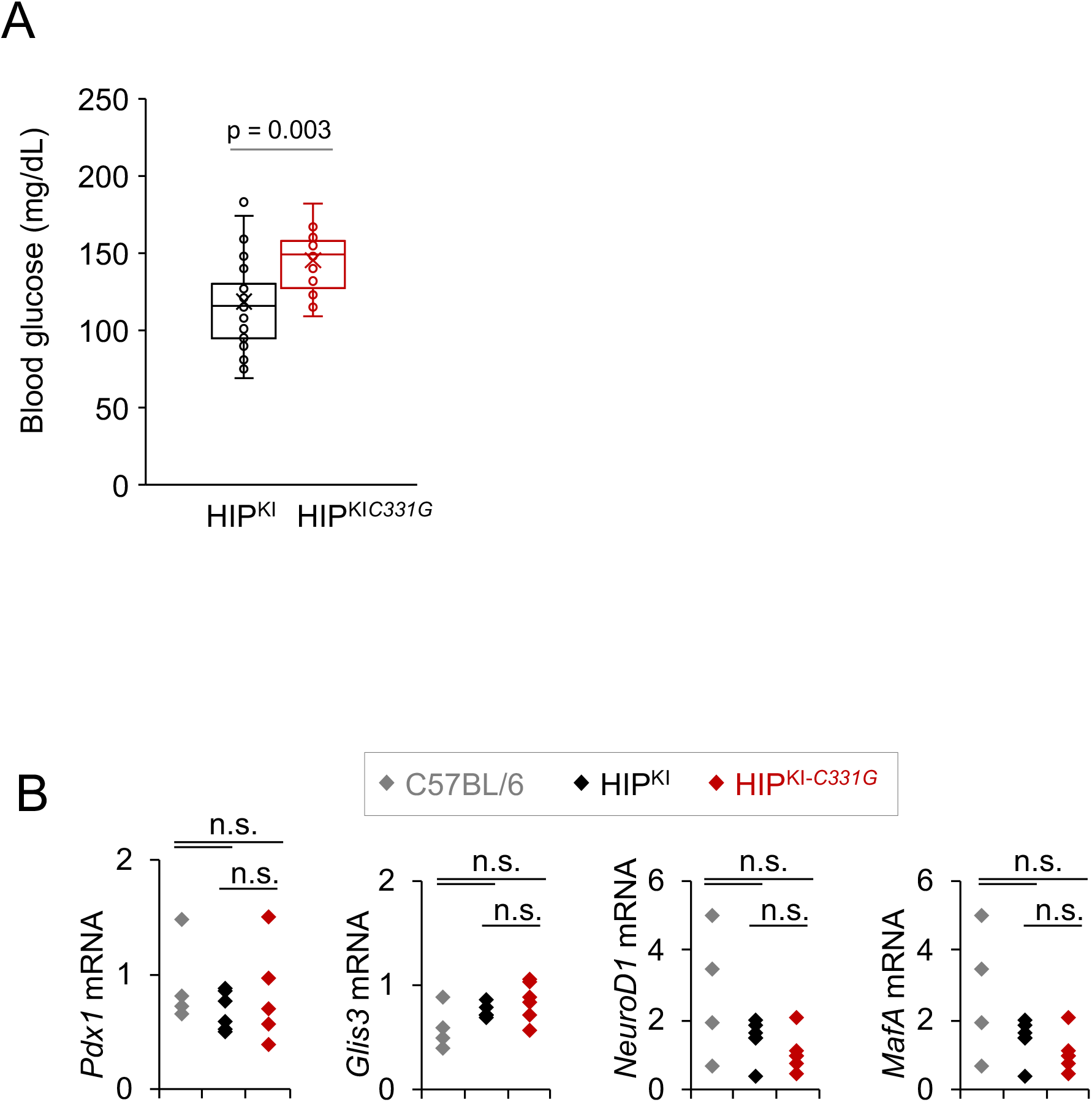
(A) Glycemia from ad libitum-fed HIP^KI^ and HIP^KI*C331G*^ mice. P values were calculated with Student’s t test. (B) Reverse transcription quantitative PCR for *Pdx1, Glis3, NeuroD1, and MafA* mRNAs from pancreatic islets from 3-5 month control C57BL/6 (n=4), HIP^KI^ (n=6) and HIP^KI-C331G^ (n=6) mice. Values were normalized to *Actb* or *Hprt* mRNAs.

**Supplementary Figure 3.**
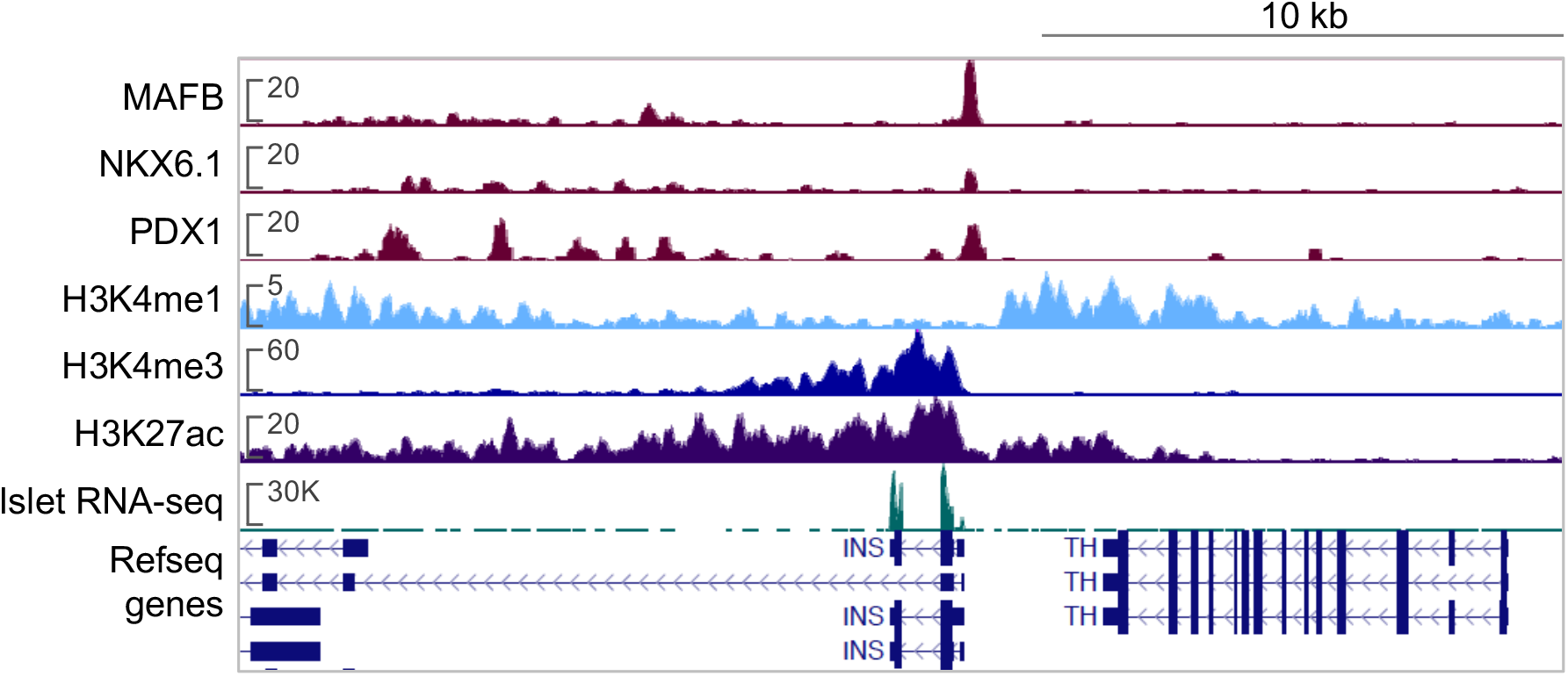
Chromatin landscape of the human *INS* locus in human pancreatic islets. ChIP-seq profiles of activating histone marks H3K4 trimethylation and H3K27 acetylation^12^. All scales represent RPKMs.

**Supplementary Figure 4.**
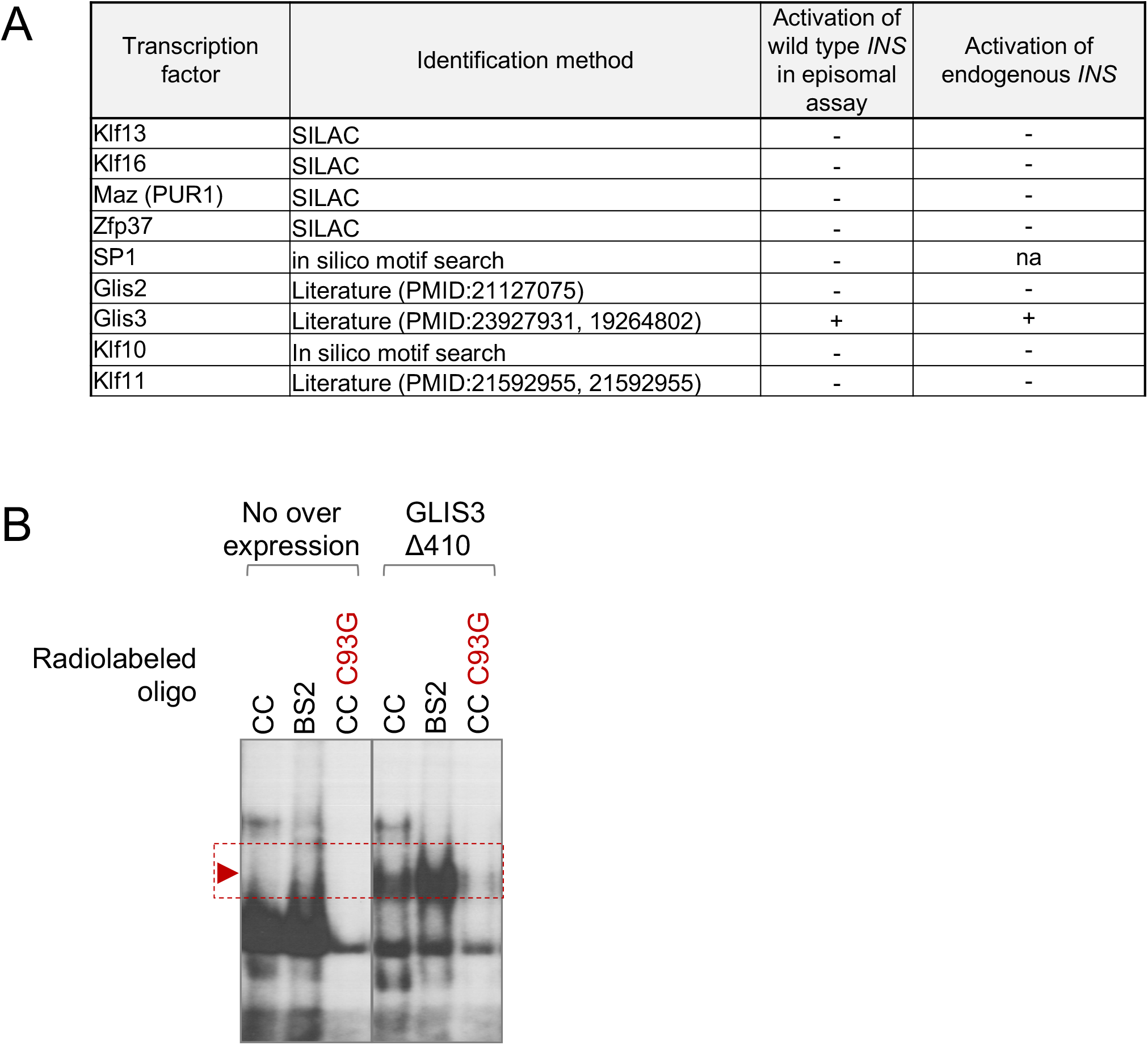
(A) Candidate DNA-binding transcription factor regulators of the *INS* gene that underlie the deleterious effects of the c.-331C>G mutation. Four transcription factors were selected based on differential binding to c.-331C vs. c.-331C>G double stranded oligonucleotides in SILAC experiments in MIN6 b cells, while others were selected based on *in silico* predicted differential binding to c.-331C vs. c.-331C>G *INS* sequences or published studies from indicated references. The summary table emphasizes that amongst these candidates only GLIS3 led to activation of the unmodified episomal insulin promoter plasmid, and activated *INS* mRNA in non-pancreatic cell lines in the presence of islet transcription factors as shown in Supplementary Figure 6. (B) Electromobility shift assays (EMSA) from HEK293T cells transfected with human GLIS3Δ410 cDNA show binding to CC element oligonucleotides and BS2, another previously reported GLIS3 recognition sequence in the INS 5’ flanking regions^29^, but not to the CC element carrying the - 331C>G mutation.

**Supplementary Figure 5.**
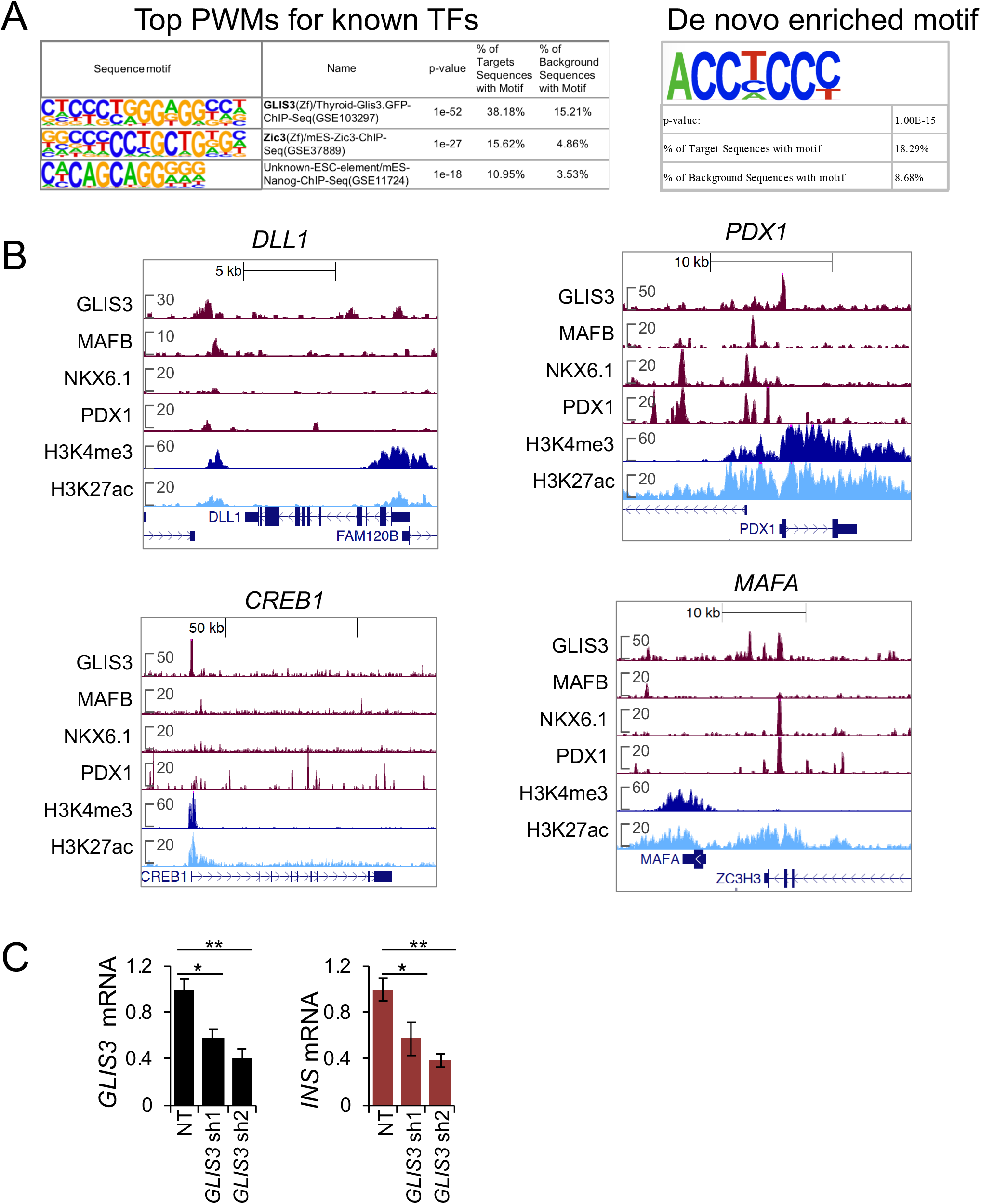
(A) The three most enriched position weight matrixes (PWMs) at GLIS3-bound regions in human islets included a previously identified GLIS3-bound PWM (left panel). Discovery of *de novo* enriched PWMs identified sequence PWMs matching known GLIS3 recognition sequences, including one matching the *INS* CC element (right panel). (B) Examples of GLIS3-bound regions in human pancreatic islets at selected loci, along with binding profiles of other human pancreatic islet transcription factors. (C) Quantitative reverse-transcription PCR of INS and GLIS3 mRNAs after lentiviral-mediated knockdown of GLIS3, using two independent shRNA sequences, in EndoCb-H1 human b-cells. Student’s t test * p < 0.05 or ** p < 0.001.

**Supplementary Figure 6.**
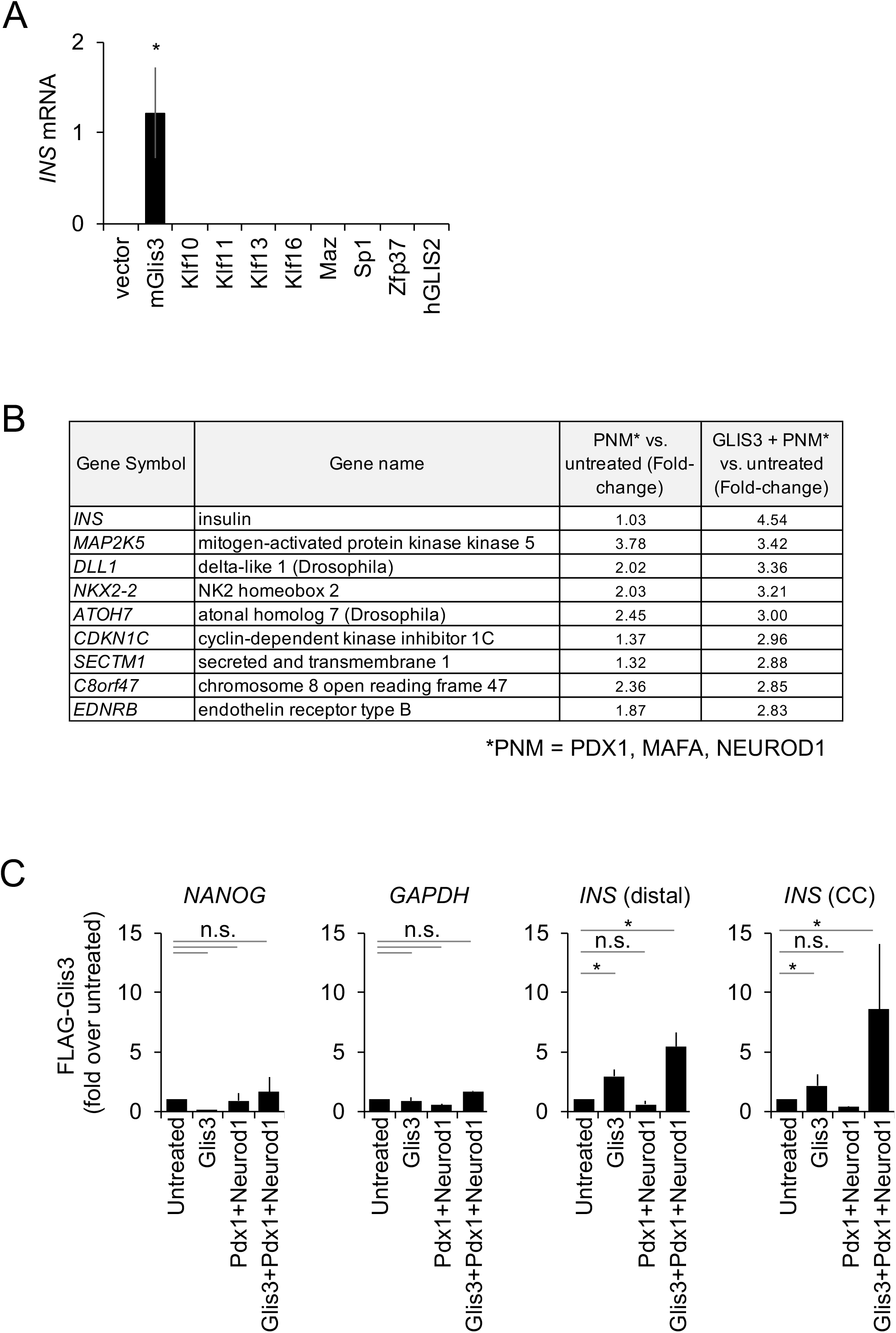
(A) GLIS3, but not other candidate transcription factors that bind CC, activates *INS* in heterologous cell types. HEK 293T cells were either left untreated (Unt) or transfected with plasmids overexpressing the indicated transcription factors along with NEUROD1 and PDX1. *INS* mRNA are *INS* to *GAPDH* mRNA ratios X 1000. Statistical significance was calculated relative to untreated samples (n = 3 independent experiments). * Student’s t test, p < 0.05. Analogous experiments were performed in the presence of MAFA, or NEUROG3 instead of NEUROD1, yielding similar results. (B) Table showing the list of genes most impacted by the overexpression of GLIS3+ PNM (PDX1, NEUROD1, MAFA) vs. PNM alone, in HEK293T cells. Cells were either left untreated or transfected with the indicated transcription factors. At 3 days post transfection RNA was quantified using microarrays. Values indicate fold-difference in expression relative to control cells. This experiment was performed with a single replicate and provides an unbiased confirmation that *INS* was the single most induced gene when GLIS3 was added to the transcription factor cocktail. (C) ChIP of FLAG-tagged GLIS3 in HEK 293T cells transfected with the indicated transcription factors. ChIP DNA was quantified by PCR and expressed as percentage of input DNA and as fold enrichment over untreated HEK293T cells. Statistical significance was calculated relative to untreated (Unt) samples using students t-test (n=3 experiments). As expected, *NANOG* promoter and *GAPDH* enhancer showed no changes between treatments.

